## Supplementary material for "UPF3A is dispensable for nonsense-mediated mRNA decay in mouse pluripotent and somatic cells": Supp Figures

### Supplementary Figures

### Supplementary Figure 1

**A**

| | Accession No. | Molecular weight (Predicted) |
|---|---|---|
| Mouse UPF3A | NP_080200.1 | 48.76 KD |
| Mouse UPF3B | NP_080849.1 | 57.07 KD |
| GFP-mUpf3a | Current study | 77.73 KD |
| GFP-mUpf3b | Current study | 86.05 KD |

Note: Molecular weights of proteins are predicted with the following website (https://www.bioinformatics.org/sms/prot_mw.html).

**B**

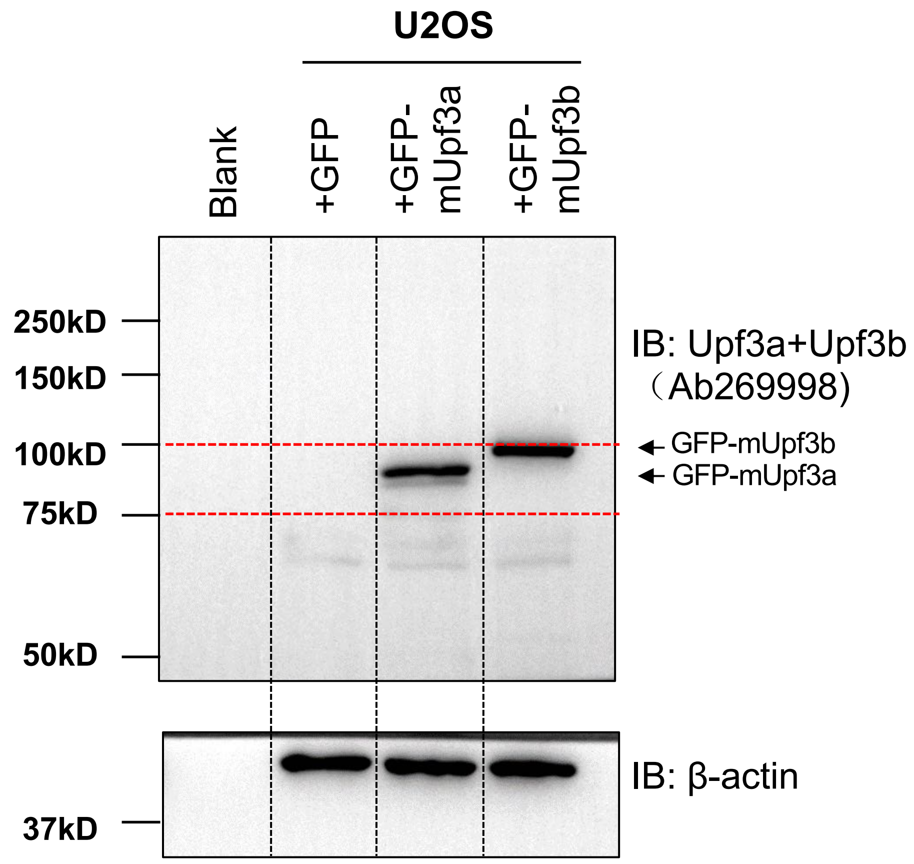


**C**

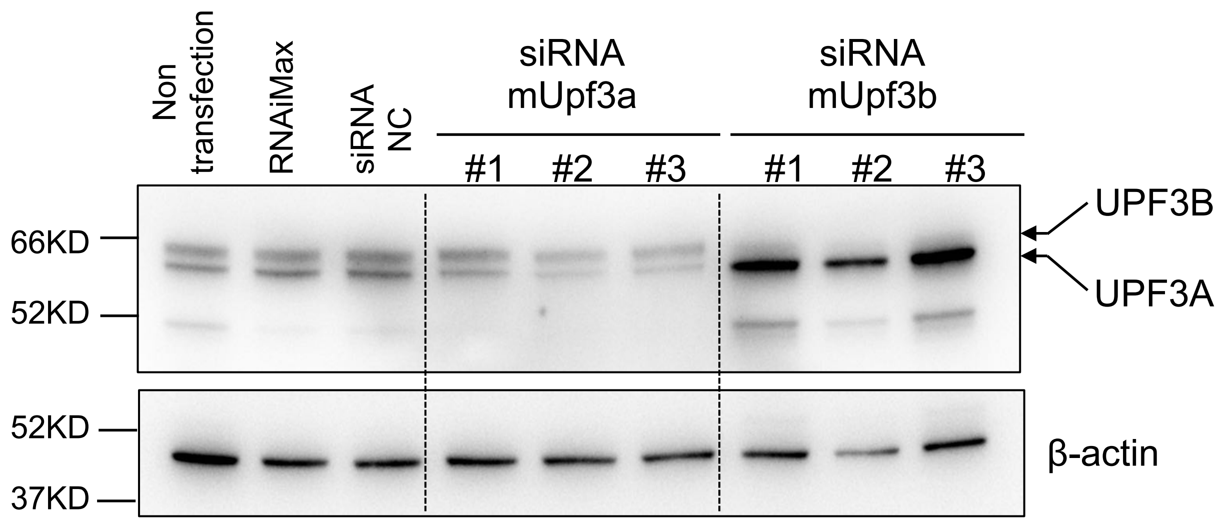

### Supplementary Figure 2

**A**

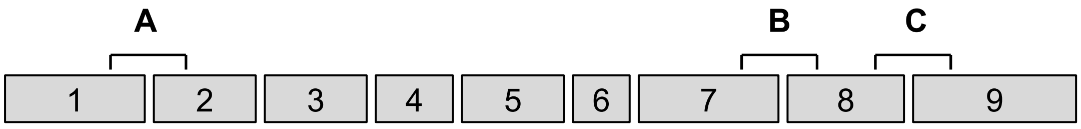


**Exons of mouse Upf3a**

| PCR fragment | Exons covered | Primer sequences |
|---|---|---|
| A | E1-E2 | F: CCCTAAGTGAGAGCGGGG<br>R: CTCTTCCAGCTGCTCTTTGG |
| B | E7-E8 | F：GTAAGAGGAAGGAGGCGGAG<br>R：TTTCTCTGTGGCCACTTCCT |
| C | E8-E9 | F: TGGAGACGAGAAGCAGGAAG<br>R: AGATCTCTTGTCCCTTGGCT |

**B**

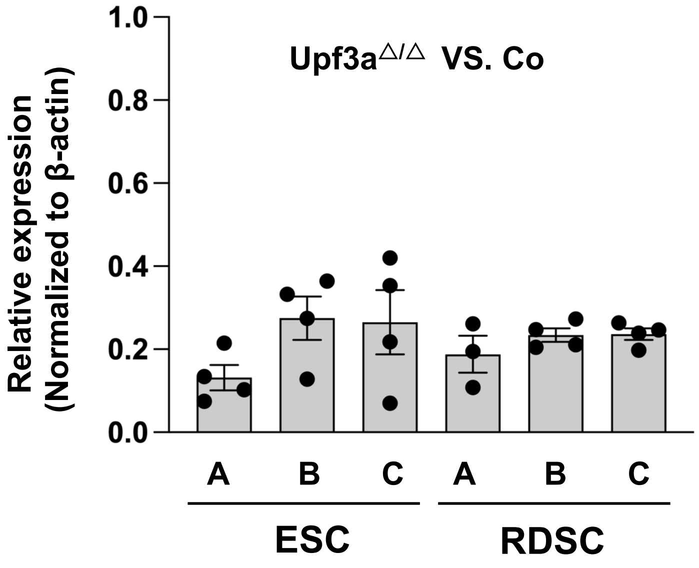

### Supplementary Figure 3

A

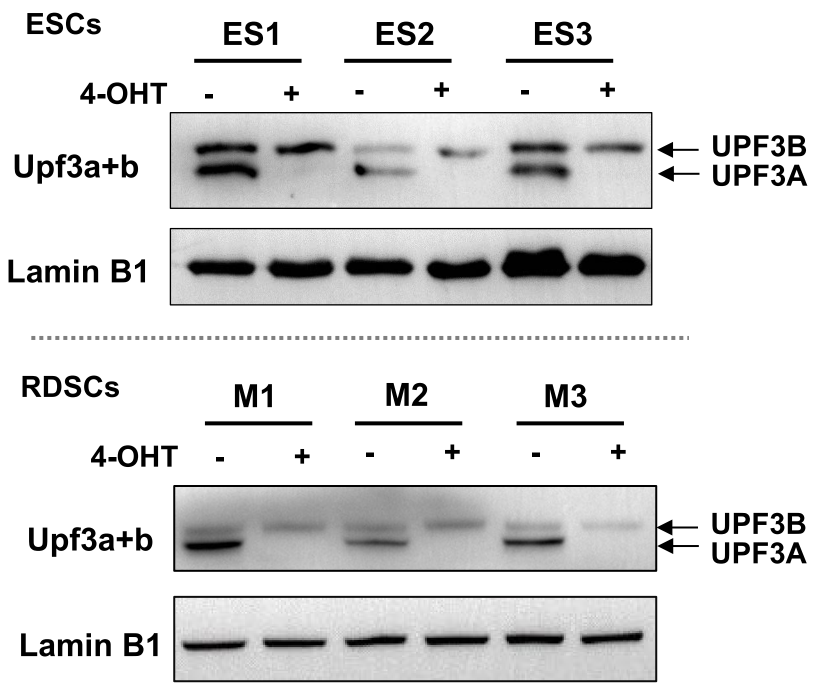


B

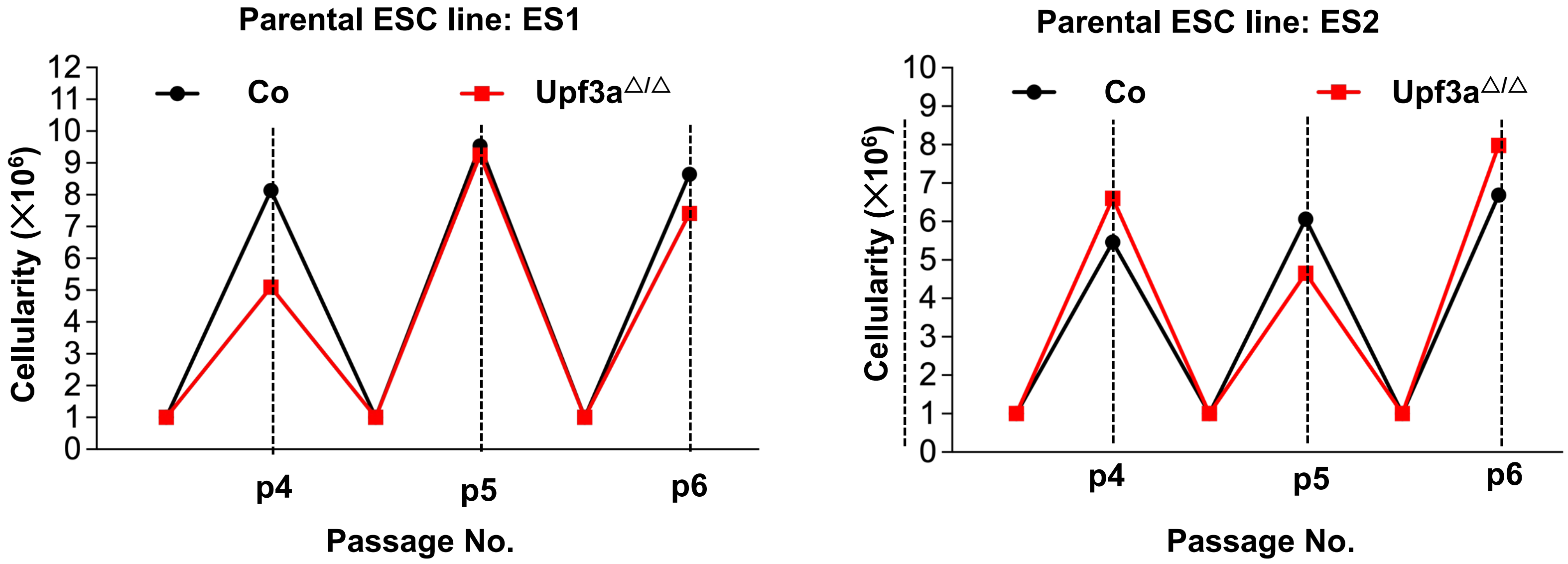

### Supplementary Figure 4

**A**

| Gene | NMD features | Reference |
|---|---|---|
| *Cdh11* | N.D | Shum *et al.* 2016, Cell |
| *Ire1* | N.D | Shum *et al.* 2016, Cell |
| *Smad5* | N.D | Shum *et al.* 2016, Cell |
| *Smad7* | N.D | Shum *et al.* 2016, Cell |
| *Snord22* | 3' UTR intron | Shum *et al.* 2016, Cell |
| *Atf4* | uORF | Weischenfeldt *et al.* 2008 , Genes Dev<br>Li *et al.* 2015, EMBO J<br>Shum *et al.* 2016, Cell |
| *Gas5* | 3' UTR intron | Weischenfeldt *et al.* 2008 , Genes Dev<br>Li *et al.* 2015, EMBO J<br>Shum et al. 2016, Cell |
| *Snhg12* | 3' UTR intron | Weischenfeldt *et al.* 2008 , Genes Dev<br>Li *et al.* 2015, EMBO J |
| *1810032 O08Rik* | PTC generated by AS | Weischenfeldt *et al.* 2008 , Genes Dev<br>Li *et al.* 2015, EMBO J |
| *Ddit3* | uORF | Weischenfeldt *et al.* 2008 , Genes Dev<br>Li *et al.* 2015, EMBO J |
| *Hnrnpl* | PTC generated by AS | Li *et al.* 2015, EMBO J |
| *Auf1* | PTC generated by AS | Li *et al.* 2015, EMBO J |
| *Smg5* | Long 3'UTR<br>uORF | Huang et al. 2011, Mol Cell<br>Li et al. 2015, EMBO J |
| *Smg6* | Long 3'UTR<br>uORF | Huang *et al.* 2011, Mol Cell<br>Li *et al.* 2015, EMBO J |
| *Smg7* | Long 3'UTR<br>uORF | Huang *et al.* 2011, Mol Cell<br>Li et al. 2015, EMBO J |
| *Upf3b* | Long 3'UTR | Huang *et al.* 2011, Mol Cell |
| *Eif4a2 (PTC)* | PTC generated by AS | Huth *et al.* 2022, Genes Dev |
| *Eif4a1 (PTC)* | PTC generated by AS | Current study |
| *Mettl23 (PTC)* | PTC generated by AS | Current study |

**Note: N.D, not determined; AS, alternative splicing.**

**B**

| Gene | NMD features | Reference |
|---|---|---|
| *Pkm2* | PTC generated by AS (exon inclusion) | Weischenfeldt *et al.* 2012 , Genome Biol |
| *Rps9* | | Weischenfeldt *et al.* 2012 , Genome Biol |
| *Eif4a2* | | McIlwain *et al.* 2010, PNAS |
| *Luc7l* | | McIlwain *et al.* 2010, PNAS |
| *Snrpb* | | McIlwain *et al.* 2010, PNAS |
| *Hnrnpa 2b1* | | McIlwain *et al.* 2010, PNAS |
| *Sfrs10* | | McIlwain *et al.* 2010, PNAS |
| *Ptbp2* | PTC generated by AS (exon exclusion) | Weischenfeldt *et al.* 2012 , Genome Biol |
| *Alkbh3* | | McIlwain *et al.* 2010, PNAS |
| *Sf1* | | McIlwain *et al.* 2010, PNAS |
| *Nfyb* | | McIlwain *et al.* 2010, PNAS |
| *Ccar1* | | McIlwain *et al.* 2010, PNAS |
| *Slc38a2* | | McIlwain *et al.* 2010, PNAS |
| *Flot1* | | McIlwain *et al.* 2010, PNAS |

### Supplementary Figure 5

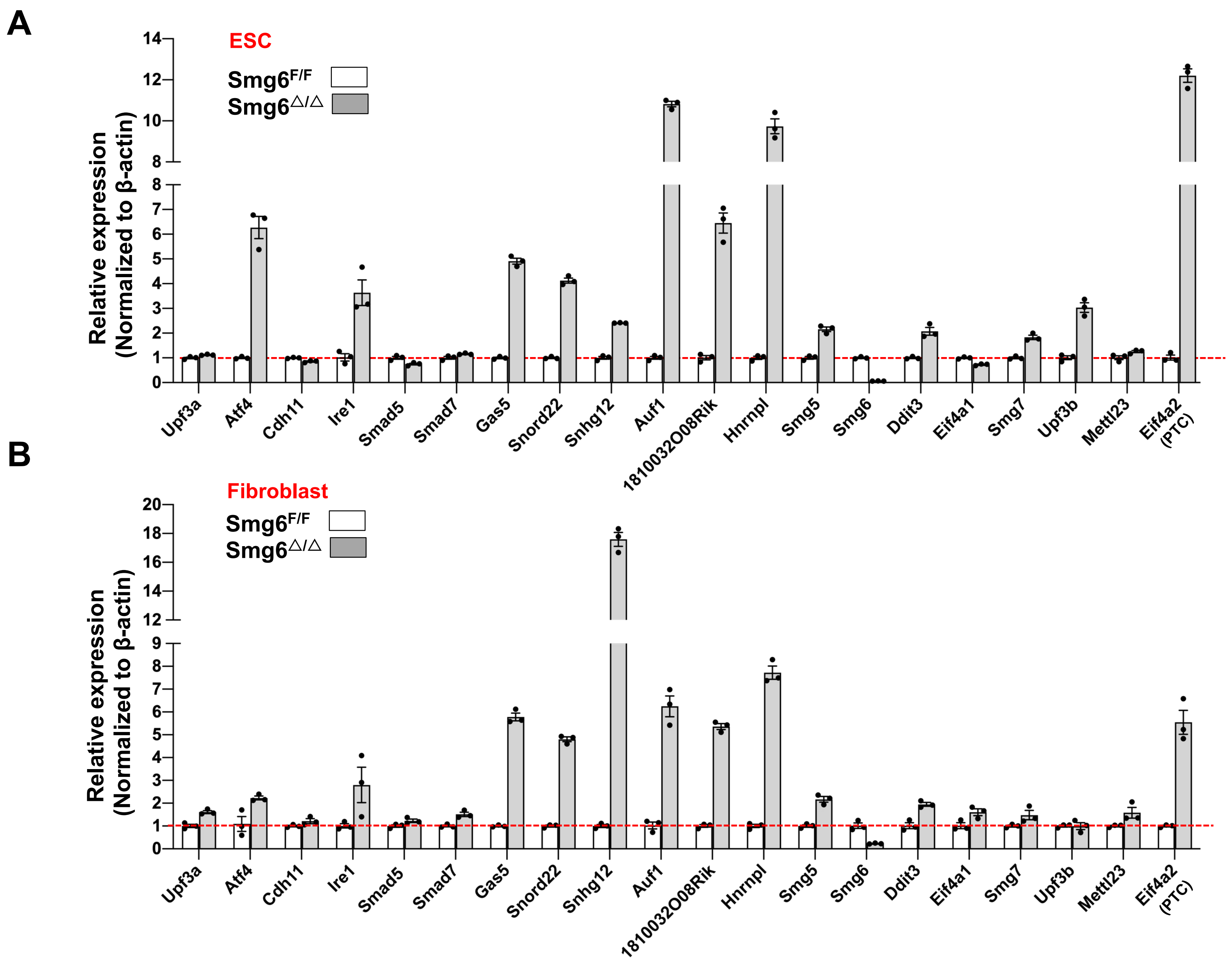

### Supplementary Figure 6

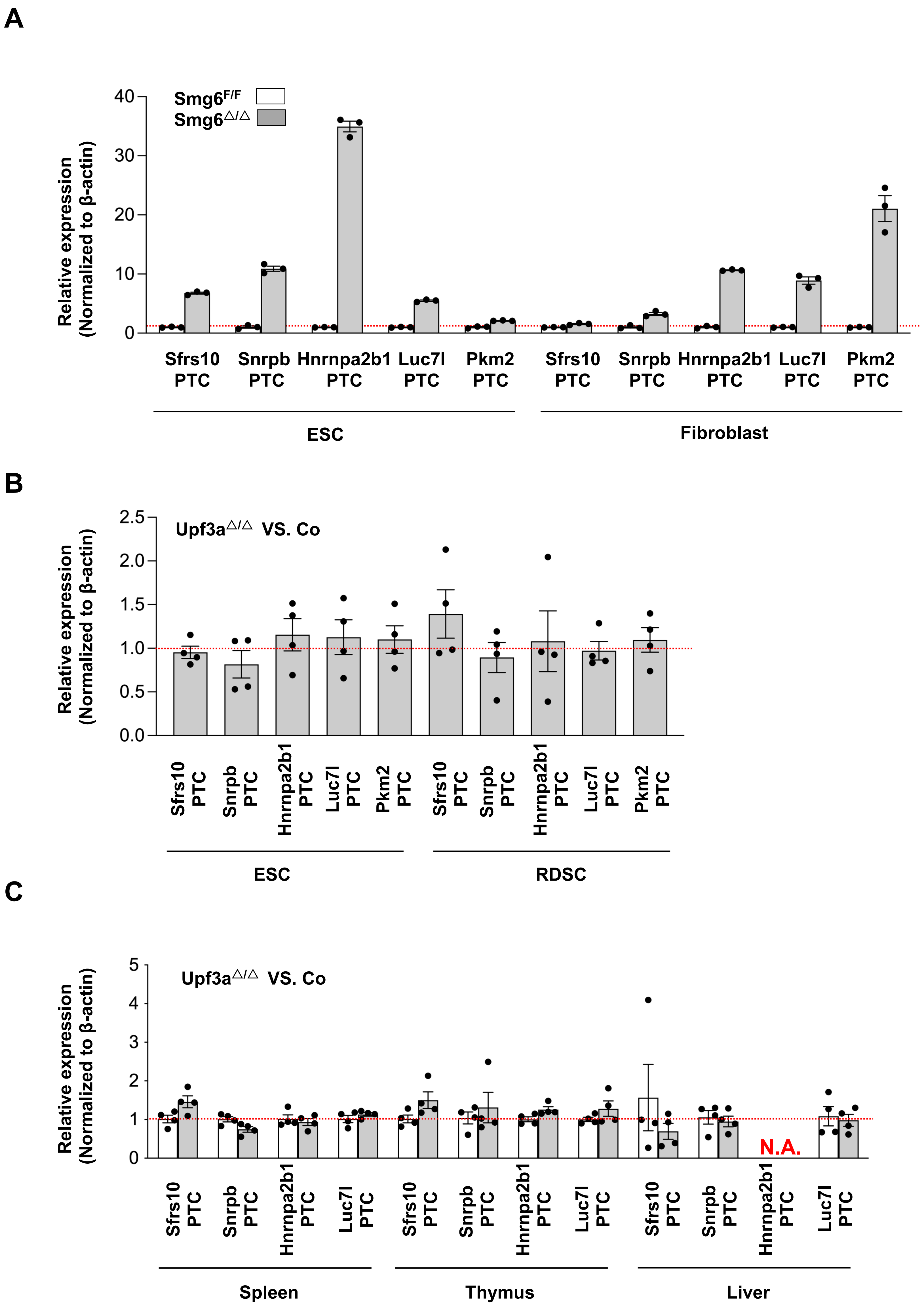

### Supplementary Figure 7

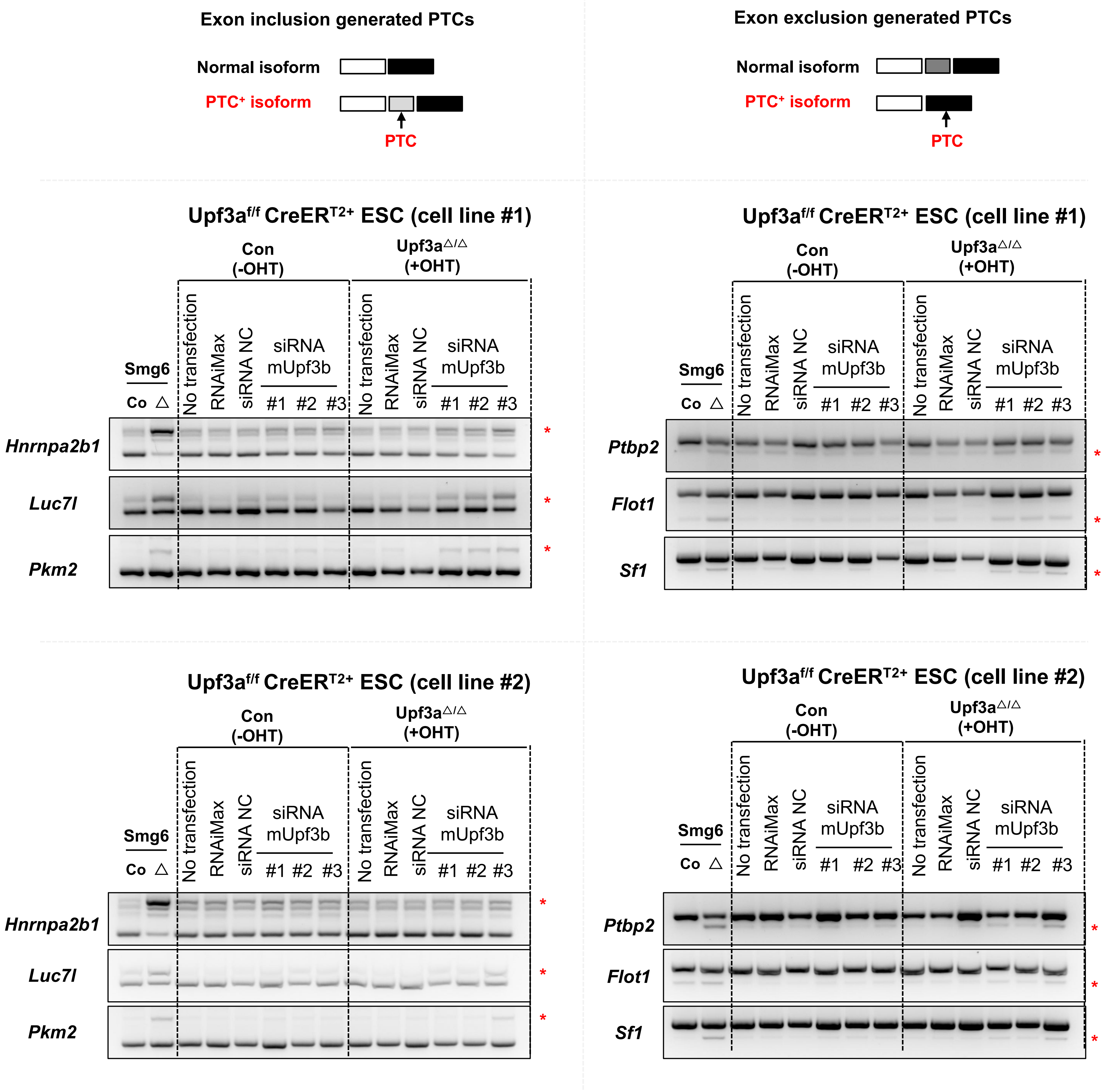

### Supplementary Figure 8

A

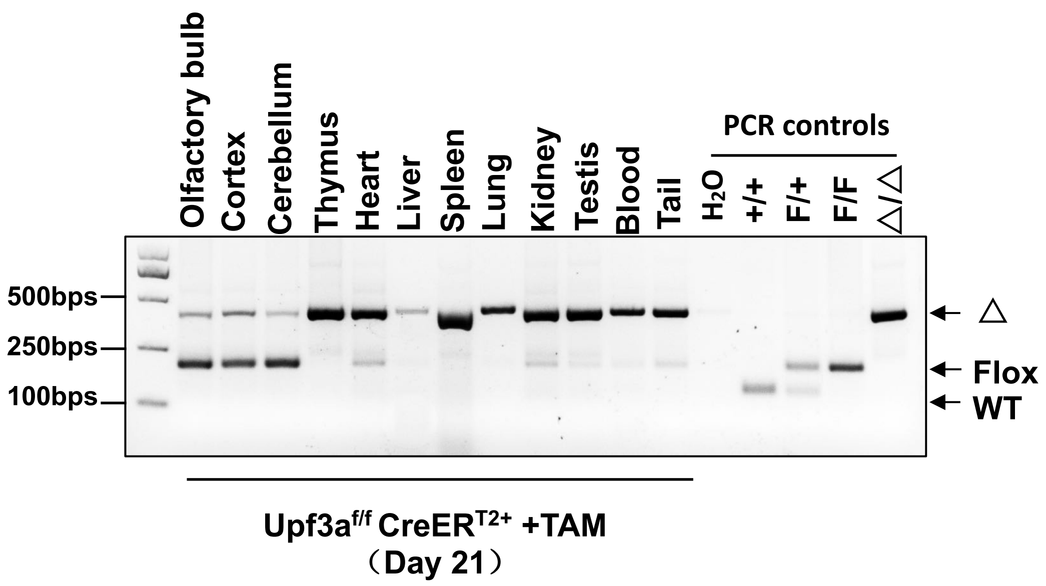


B

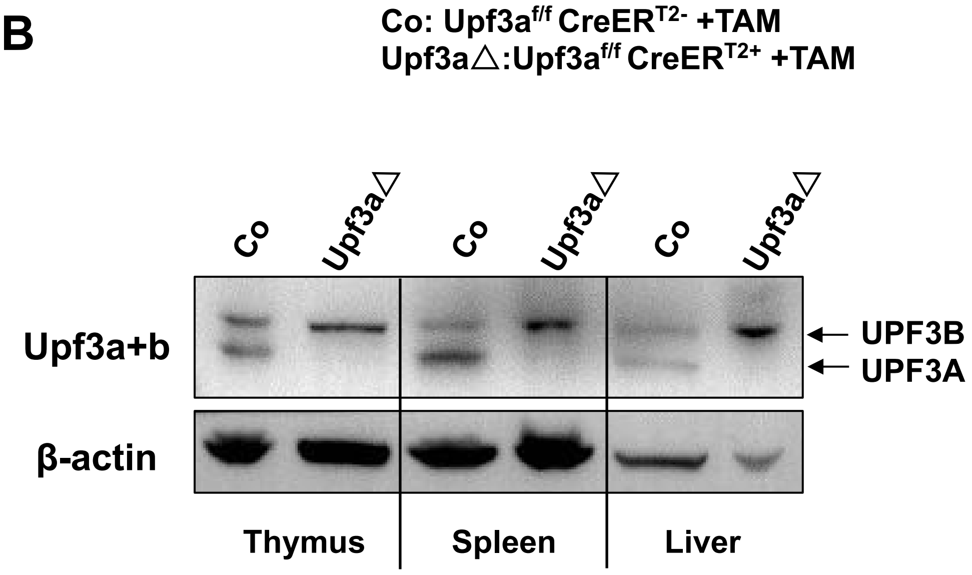

### Supplementary Figure 9

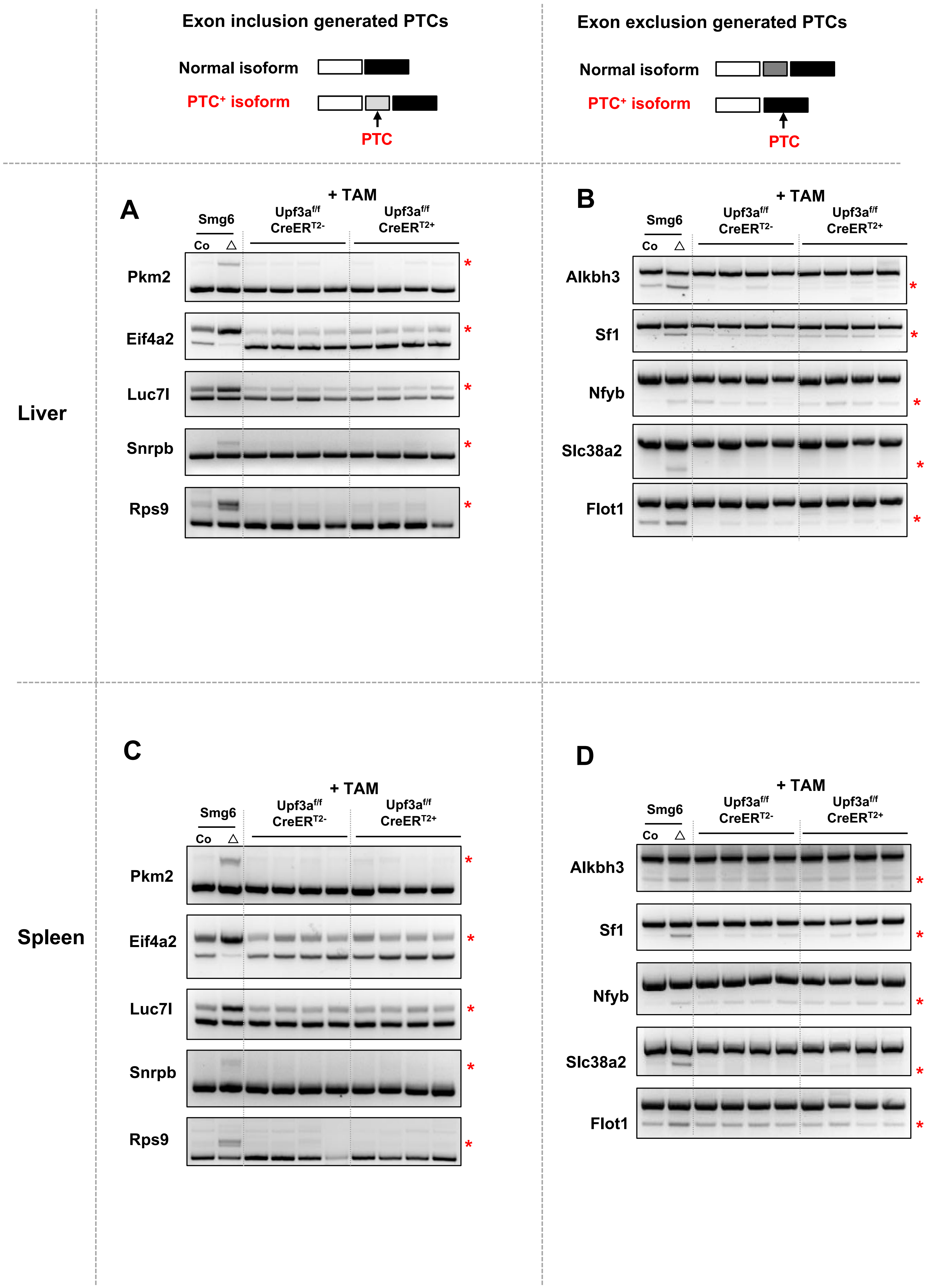
